# Evaluation of correlates of protection against influenza A(H3N2) and A(H1N1)pdm09 infection: Applications to the hospitalized patient population

**DOI:** 10.1101/416628

**Authors:** Joshua G. Petrie, Emily T. Martin, Rachel Truscon, Emileigh Johnson, Caroline K. Cheng, EJ McSpadden, Ryan E. Malosh, Adam S. Lauring, Lois E. Lamerato, Maryna C. Eichelberger, Jill M. Ferdinands, Arnold S. Monto

## Abstract

**Background:** Influenza vaccines are important for prevention of influenza-associated hospitalization. Assessments of serologic correlates of protection can support interpretation of influenza vaccine effectiveness evaluations in hospitalized populations.

**Methods:** Serum specimens collected at admission from adults hospitalized for treatment of acute respiratory illnesses during two influenza seasons were tested in hemagglutination-inhibition (HAI) and neuraminidase-inhibition (NAI) assays. We evaluated the suitability of these specimens as proxies for pre-infection immune status, and measured associations between antibody titers and influenza vaccination and infection

**Results:** Specimens were collected within 3 days of illness onset from 65% of participants; geometric mean titers (GMTs) did not vary by day of collection. In both seasons, vaccinated participants had higher HAI and NAI GMTs than unvaccinated participants. HAI titers against the 2014-2015 A(H3N2) vaccine strain did not correlate with protection from infection with antigenically-drifted A(H3N2) viruses that circulated that season. In contrast, higher HAI titers against the A(H1N1)pdm09 vaccine strain were associated with reduced odds of A(H1N1)pdm09 infection in 2015-2016.

**Conclusions:** Serum collected after hospital admission can be used to assess correlates of protection against influenza infection. Broader implementation of similar studies would provide an opportunity to understand the successes and shortcomings of current influenza vaccines.

---

Influenza infections are responsible for substantial numbers of hospitalizations and deaths each year [1]. Therefore, in the US, all individuals ≥6 months of age are recommended to receive annual influenza vaccination to prevent clinical influenza and the severe complications associated with influenza illness [2]. Continuous evolution of the influenza virus necessitates annual assessment of the strains composing influenza vaccines with vaccine strain selection occurring well in advance of the subsequent influenza season [3]. Because influenza vaccine effectiveness (VE) can vary, particularly by the antigenic match between vaccine strain and circulating epidemic viruses, annual VE assessments are necessary to inform policy makers, health care providers, and the public of the effectiveness of the influenza vaccine.

As a result, influenza VE is now evaluated annually in many countries with the majority of studies carried out in outpatient settings [4–6]. A number of issues with current influenza vaccines have now become clear, many of which are influenza type and subtype specific. On average, influenza vaccines have been only approximately 30% effective in preventing influenza A(H3N2) infection, and were ineffective in preventing antigenically drifted viruses during the 2014-2015 season [7–9]. Reasons for low VE against A(H3N2) are not completely understood, but given consistently lower VE, it is likely that factors contribute in addition to antigenic match between the hemagglutinin of vaccine and circulating viruses [10]. A potentially related issue is that reduced VE has been frequently demonstrated among individuals vaccinated in consecutive seasons compared to those vaccinated only in the current season [11].

Immunologic evaluations are needed to help explain the factors underlying low VE, improve vaccine strain selection, and guide novel vaccine development [12]. Unfortunately, in outpatient settings, it is difficult to integrate immunologic evaluations because subject participation is generally limited to a short interaction during acute illness. VE studies are increasingly being carried out among inpatient populations [13–16]. Hospitalized patients regularly have blood collected at hospital admission for clinical purposes that could be used to support VE estimates from hospital-based studies. It is unclear how well antibody titers measured in these specimens reflect pre-infection immune status, because the antibody response to infection can begin within days of illness onset.

In this study we aimed to determine if antibody titers measured in serum specimens collected from hospitalized patients during acute influenza illness accurately reflect pre-infection humoral immunity, and if so, to determine how well influenza vaccine specific antibodies correlated with odds of infection. The study was performed during the 2014-2015 and 2015-2016 influenza seasons. These two seasons differed in the predominant circulating strains and the resulting VE estimates. In 2014-15, a drifted influenza A(H3N2) virus belonging to the 3C.2a genetic group predominated in the United States[9]. The vaccine was ineffective in ambulatory populations against drifted viruses, but moderately effective against other circulating genetic groups that were more closely related to the vaccine virus [9]. In contrast, we estimated modest, but statistically significant VE against influenza A(H3N2) in this hospitalized population [13]. The following year, 2015-2016, influenza A(H1N1)pdm09 viruses predominated, and VE was 45% against influenza A(H1N1)pdm09 in the ambulatory setting [17].

## METHODS

During the 2014-2015 and 2015-2016 influenza seasons adult (≥18 years) patients hospitalized for treatment of acute respiratory illnesses at the University of Michigan Hospital in Ann Arbor and the Henry Ford (HF) Hospital in Detroit were prospectively enrolled in a case-test negative design study of influenza vaccine effectiveness as previously described [13]. All patients were enrolled ≤10 days from illness onset during the period of influenza circulation each year (November-March 2014-2015; January-April 2015-2016). Participants completed an enrollment interview and had throat and nasal swab specimens collected and combined for influenza identification. When available, clinical serum specimens collected as early as possible after hospital admission were retrieved; all specimens were collected ≤10 days from illness onset based on the enrollment case definition. Serum specimens were available only for patients enrolled from University of Michigan Hospital during the 2014-2015 season, but were available for patients enrolled from both hospitals during the 2015-2016 season. The institutional review boards at both participating health systems reviewed and approved the study.

### Participant Data

Enrolled patients were interviewed to collect information on demographic characteristics (age, sex, race/ethnicity), illness onset date, self-reported influenza vaccination status, and frailty (instrument fully described in Supplement). Health System Electronic Medical Records were reviewed to document evidence of health conditions present during the year before enrollment to calculate the Charlson comorbidity index. Influenza vaccination in each season was based on documented evidence of vaccine receipt from the Electronic Medical Record or statewide immunization registry or from self-report of vaccine receipt at enrollment with plausible location and timing of vaccination (plausible self-report).

### Laboratory Assays

Respiratory specimens were tested for influenza by RT-PCR in the investigators’ laboratory at the University of Michigan School of Public Health using primers, probes and testing protocols developed by the Influenza Division of the Centers for Disease Control and Prevention (CDC) and provided by the CDC’s International Reagent Resource.

Serum specimens collected from patients enrolled during the 2014-2015 influenza season were tested in hemagglutination-inhibition (HAI) assays using monovalent inactivated influenza vaccine subunit material (provided by Sanofi-Pasteur) representing the A/Texas/50/2012 (H3N2) virus present in the 2014-2015 Northern Hemisphere influenza vaccine and the A/Hong Kong /4801/2014 (H3N2) virus present in the 2016-2017 Northern Hemisphere influenza vaccine, representing the dominant variant 3C.2a genetic group that circulated in 2014-2015. Serum specimens collected from patients enrolled during the 2015-2016 influenza season were similarly tested by HAI using monovalent inactivated influenza vaccine subunit material representing the A/California/7/2009 (H1N1)pdm09 virus present in the 2015-2016 Northern Hemisphere influenza vaccine. Serum specimens collected in the 2014-2015 and 2015-2016 seasons were also tested by neuraminidase inhibition (NAI) assay using reassortant influenza viruses with the NA of the 2014-2015 A(H3N2) and 2015-2016 A(H1N1)pdm09 vaccine strains, respectively, and a mismatched HA (H6 subtype) to avoid interference by HA-specific antibodies[18]. Serologic testing was performed in the investigators’ laboratory at the University of Michigan School of Public Health (methods for each assay are fully described in Supplement).

### Statistical Analysis

Subjects were characterized by demographics, health status, influenza vaccination status, and influenza infection status. HAI and NAI titers were calculated as the reciprocal (e.g., 40) of the highest dilution of sera (e.g., 1:40) that inhibited HA or NA activity. Titers were log-base 2 transformed, and the mean of the transformed values was calculated and then exponentiated to obtain the geometric mean titer (GMT). GMTs were compared by subject characteristics including health, current and prior vaccination, and infection status using unbalanced ANOVA F-tests or Wilcoxon Rank Sum tests.

To evaluate the assumption that antibody titers measured in specimens during acute illness reflect pre-infection titers rather than a de novo response to infection, we estimated the association between log-base-2 antibody titer and days from illness onset to specimen collection among individuals with laboratory-confirmed influenza infection using linear regression models. If measured antibody titers reflect a de novo response, those collected later in the illness would be expected to have higher antibody titers, which would be reflected in a positive slope estimated by the regression. Similarly, we evaluated the possibility that influenza negative controls, particularly those enrolled later in the season, could have been infected before the time of enrollment. This was accomplished by estimating the association between log-base-2 antibody titer and day of illness onset within each season among influenza negative controls using linear regression models. If a substantial proportion of control subjects were infected at some time prior to enrollment, those enrolled later in the season would be expected to have higher antibody titers, which would be reflected in a positive slope estimated by the regression.

We estimated the change in odds of influenza infection (Odds Ratio [OR]) associated with a 2-fold increase in preseason HAI and NAI titers modeled as continuous, log-base 2 transformed predictors in logistic regression models for each influenza season. We specified models to assess the effect of each titer individually, to assess the independent effect of each titer holding other measured titers constant (partial adjustment), and to assess the effect of each titer after adjustment for other measured titers and covariates identified a priori as potential confounding variables associated with both antibody titer and odds of infection (full adjustment). The following covariates were included in fully adjusted models: vaccination status, gender, age category (18-49, 50-64, ≥65 years), frailty score, and Charlson comorbidity index category (0, 1, 2, ≥3).

All statistical analyses were performed using SAS (release 9.3; SAS Institute); figures were prepared using R software (version 3.1.0). A *P*-value of <.05 indicated statistical significance.

## RESULTS

During the 2014-2015 influenza season, 624 patients with acute respiratory illness were enrolled in the VE study. Of these, 341 were enrolled at the University of Michigan Hospital and serum specimens were retrieved for 315 (92%) of these patients. Serum specimens were retrieved for 339 (77%) of all 441 patients enrolled from both hospitals during the 2015-2016 influenza season. At the University of Michigan site, subject characteristics of those with retrieved specimens (2014-2015: 92%; 2015-2016: 95%) were similar to those of all patients enrolled at the University of Michigan Hospital for both seasons (Supplemental Tables 1 and 2). However, subjects with specimens retrieved from the Henry Ford Hospital during the 2015-2016 season (52%) were more likely to be white (28% vs 13%, *P*=0.03) and have higher Charlson comorbidity index (3+: 53% vs 38%, *P*=0.02) compared to those subjects for whom specimens were unable to be retrieved (Supplemental Table 2). The characteristics of the subjects with specimens retrieved during each season are presented in Tables 1 and 2, respectively.

**Table 1.** Geometric mean antibody titers among adults hospitalized for acute respiratory illness during the 2014-2015 influenza season by subject characteristics.

| Characteristic | Participants<br>No. (%) | A/Texas/50/2012 (H3N2)<br>HAI GMT (95% CI) | P-value<sup>a</sup> | A/Texas/50/2012 (H3N2)<br>NAI GMT (95% CI) | P-value<sup>a</sup> | A/Hong Kong/4801/2014 (H3N2)<br>HAI GMT (95% CI) | P-value<sup>a</sup> |
| --- | --- | --- | --- | --- | --- | --- | --- |
| Overall | 315 (100.0) | 38.8 (33.5, 44.8) | -- | 124.5 (109.3, 141.9) | -- | 10.4 (9.2, 11.8) | -- |
| Study hospital |  |  |  |  |  |  |  |
| University of Michigan, Ann Arbor | 315 (100.0) | 38.8 (33.5, 44.8) | -- | 124.5 (109.3, 141.9) | -- | 10.4 (9.2, 11.8) | -- |
| Sex |  |  |  |  |  |  |  |
| Female | 155 (49.2) | 42.4 (34.5, 52.1) | 0.238 | 123.4 (102.1, 149.3) | 0.900 | 11.1 (9.2, 13.3) | 0.332 |
| Male | 160 (50.8) | 35.6 (29.0, 43.7) |  | 125.5 (104.9, 150.2) |  | 9.8 (8.4, 11.5) |  |
| Age category (y) |  |  |  |  |  |  |  |
| 18-49 | 84 (26.7) | 32.5 (23.9, 44.3) | 0.126 | 92.0 (70.4, 120.4) | 0.021 | 9.9 (7.8, 12.6) | 0.604 |
| 50-64 | 98 (31.1) | 35.7 (27.6, 46.3) |  | 134.1 (106.0, 169.6) |  | 9.9 (7.9, 12.3) |  |
| ≥65 | 133 (42.2) | 46.0 (37.4, 56.7) |  | 142.7 (118.3, 172.1) |  | 11.2 (9.3, 13.5) |  |
| Race/ethnicity<sup>b</sup> |  |  |  |  |  |  |  |
| White (not Hispanic) | 242 (77.3) | 35.4 (29.9, 41.8) | 0.062 | 114.8 (98.9, 133.1) | 0.082 | 9.9 (8.6, 11.3) | 0.322 |
| Black | 43 (13.7) | 52.6 (36.7, 75.4) |  | 173.4 (117.5, 256.0) |  | 12.7 (8.8, 18.5) |  |
| Other | 28 (8.9) | 55.2 (34.8, 87.5) |  | 144.9 (103.1, 203.7) |  | 11.6 (7.9, 17.0) |  |
| Frailty score (range 0-5) |  |  |  |  |  |  |  |
| 0-2 | 226 (71.8) | 37.0 (31.4, 43.8) | 0.324 | 121.8 (105.0, 141.3) | 0.597 | 9.6 (8.4, 11.0) | 0.043 |
| 3-5 | 89 (28.3) | 43.6 (32.5, 58.4) |  | 131.7 (100.8, 172.1) |  | 12.7 (9.9, 16.4) |  |
| Charlson comorbidity index category |  |  |  |  |  |  |  |
| 0 | 33 (10.5) | 30.4 (18.2, 50.8) | 0.172 | 129.7 (81.0, 207.7) | 0.730 | 10.7 (7.1, 16.1) | 0.844 |
| 1 | 71 (22.5) | 33.2 (24.3, 45.4) |  | 110.4 (85.2, 143.2) |  | 9.4 (7.6, 11.7) |  |
| 2 | 53 (16.8) | 52.6 (37.0, 74.8) |  | 118.4 (84.9, 165.1) |  | 11.1 (7.9, 15.7) |  |
| 3+ | 158 (50.2) | 39.5 (32.5, 48.0) |  | 132.5 (110.7, 158.6) |  | 10.6 (9.0, 12.6) |  |
| Illness onset to blood collection |  |  |  |  |  |  |  |
| 0-3 days | 206 (65.4) | 42.8 (35.8, 51.1) | 0.184 | 128.1 (108.8, 150.8) | 0.584 | 11.0 (9.4, 12.8) | 0.489 |
| 4-6 days | 75 (23.8) | 32.9 (24.2, 44.8) |  | 125.8 (95.3, 166.2) |  | 9.6 (7.7, 12.1) |  |
| 7-10 days | 34 (10.8) | 30.7 (20.2, 46.6) |  | 102.2 (73.2, 142.7) |  | 9.0 (6.4, 12.7) |  |
| Influenza vaccination status |  |  |  |  |  |  |  |
| Vaccinated | 237 (75.2) | 47.4 (40.5, 55.5) | <0.001 | 153.1 (133.2, 176.1) | <0.001 | 11.7 (10.1, 13.6) | <0.001 |
| Unvaccinated | 78 (24.8) | 21.1 (15.6, 28.5) |  | 66.4 (50.7, 86.9) |  | 7.3 (6.0, 8.8) |  |
| Influenza Status |  |  |  |  |  |  |  |
| A(H3N2) Positive | 43 (13.7) | 29.4 (19.7, 44.0) | 0.139 | 98.7 (71.1, 136.9) | 0.165 | 7.9 (6.3, 9.7) | 0.068 |
| Negative | 272 (86.3) | 40.5 (34.7, 47.3) |  | 129.2 (112.1, 148.8) |  | 10.9 (9.5, 12.5) |  |

Abbreviations: 95% CI, 95% confidence interval; GMT, geometric mean titer; HAI, hemagglutination-inhibition; NAI, neuraminidase-inhibition.
<sup>a</sup> P-values are calculated from unbalanced ANOVA F-test on the original log titers. <sup>b</sup> 2 participants had missing Race/ethnicity data

**Table 2.** Geometric mean antibody titers among adults hospitalized for acute respiratory illness during the 2015-2016 influenza season by subject characteristics.

| Characteristic | Participants<br>No. (%) | A/California/7/2009 (H1N1)pdm09<br>HAI GMT (95% CI) | P-value<sup>a</sup> | A/California/7/2009 (H1N1)pdm09<br>NAI GMT (95% CI) | P-value<sup>a</sup> |
| --- | --- | --- | --- | --- | --- |
| Overall | 339 (100.0) | 55.9 (47.9, 65.3) | -- | 50.5 (42.5, 59.9) | -- |
| Study hospital |  |  |  |  |  |
| University of Michigan, Ann Arbor | 243 (71.7) | 56.8 (47.5, 68.0) | 0.754 | 44.8 (36.6, 55.0) | 0.030 |
| Henry Ford, Detroit | 96 (28.3) | 53.8 (39.8, 72.8) |  | 68.3 (50.1, 93.0) |  |
| Sex |  |  |  |  |  |
| Female | 187 (55.2) | 53.6 (43.9, 65.4) | 0.550 | 44.2 (35.5, 55.0) | 0.091 |
| Male | 152 (44.8) | 58.9 (46.3, 75.1) |  | 59.5 (45.4, 77.9) |  |
| Age category (y) |  |  |  |  |  |
| 18-49 | 106 (31.3) | 64.5 (47.1, 88.2) | 0.442 | 35.8 (26.2, 48.8) | <0.001 |
| 50-64 | 122 (36.0) | 54.4 (41.8, 70.7) |  | 43.8 (32.8, 58.4) |  |
| ≥65 | 111 (32.7) | 50.4 (40.3, 63.0) |  | 82.0 (62.4, 107.7) |  |
| Race/ethnicity<sup>b</sup> |  |  |  |  |  |
| White (not Hispanic) | 198 (59.3) | 56.0 (46.5, 67.4) | 0.977 | 45.9 (36.8, 57.1) | 0.433 |
| Black | 95 (28.4) | 56.8 (40.6, 79.4) |  | 58.5 (41.7, 82.0) |  |
| Other | 41 (12.3) | 59.0 (37.3, 93.3) |  | 56.1 (34.9, 90.2) |  |
| Frailty score (range 0-5) |  |  |  |  |  |
| 0-2 | 232 (68.4) | 52.5 (43.7, 63.0) | 0.236 | 48.6 (39.3, 60.0) | 0.513 |
| 3-5 | 107 (31.6) | 64.2 (48.2, 85.4) |  | 54.9 (41.0, 73.7) |  |
| Charlson comorbidity index category |  |  |  |  |  |
| 0 | 41 (12.1) | 36.8 (23.1, 58.5) | 0.230 | 29.0 (19.3, 43.5) | 0.014 |
| 1 | 55 (16.2) | 53.4 (37.0, 77.2) |  | 35.7 (24.0, 53.1) |  |
| 2 | 45 (13.3) | 57.9 (37.0, 90.6) |  | 52.8 (32.7, 85.3) |  |
| 3+ | 198 (58.4) | 61.3 (50.3, 74.8) |  | 61.7 (49.1, 77.7) |  |
| Illness onset to blood collection <sup>c</sup> |  |  |  |  |  |
| 0-3 days | 194 (58.6) | 58.8 (47.2, 73.3) | 0.538 | 53.8 (42.5, 68.1) | 0.586 |
| 4-6 days | 97 (29.3) | 49.9 (38.5, 64.6) |  | 43.9 (32.7, 59.0) |  |
| 7-10 days | 40 (12.1) | 47.6 (32.4, 69.9) |  | 47.6 (28.8, 78.4) |  |
| Influenza vaccination status |  |  |  |  |  |
| Vaccinated | 237 (69.9) | 78.8 (66.8, 93.1) | <0.001 | 65.6 (53.9, 79.7) | <0.001 |
| Unvaccinated | 102 (30.1) | 25.2 (19.0, 33.5) |  | 27.5 (20.1, 37.7) |  |
| Influenza Status |  |  |  |  |  |
| A(H1N1)pdm09 Positive | 64 (18.9) | 23.5 (17.2, 32.3) | <0.001 | 23.3 (16.3, 33.3) | <0.001 |
| Negative | 275 (81.1) | 68.4 (57.9, 80.9) |  | 60.5 (50.1, 73.0) |  |

Abbreviations: 95% CI, 95% confidence interval; GMT, geometric mean titer; HAI, hemagglutination-inhibition; NAI, neuraminidase-inhibition.
<sup>a</sup> P-values are calculated from unbalanced ANOVA F-test. <sup>b</sup> 5 participants had missing race/ethnicity data <sup>c</sup> 8 participants had missing time between onset and blood collection

Blood specimens were collected 0-10 days after illness onset; specimens were collected within 3 days of illness from 65% of patients and within 6 days from 89%. Overall, GMTs did not significantly vary by the time of specimen collection. The effect of time of specimen relative to illness onset on antibody titers was further explored among influenza positive cases, and we did not observe an association between timing of blood collection and HAI or NAI antibody titers against viruses similar to those that circulated in each season (Figure 1). In addition, the variance of the measured titers did not appear to be associated with the timing of blood collection. This supports our assumption that measured antibody titers reflect pre-infection susceptibility rather than a de novo response to infection. Among influenza negative subjects, we also did not observe an association between day of illness onset and HAI or NAI antibody titers (or their variance) against viruses similar to those that circulated in each season (Supplemental Figure 1). This suggests that a there was not substantial misclassification of case-control status based on calendar time of enrollment.

**Figure 1.**
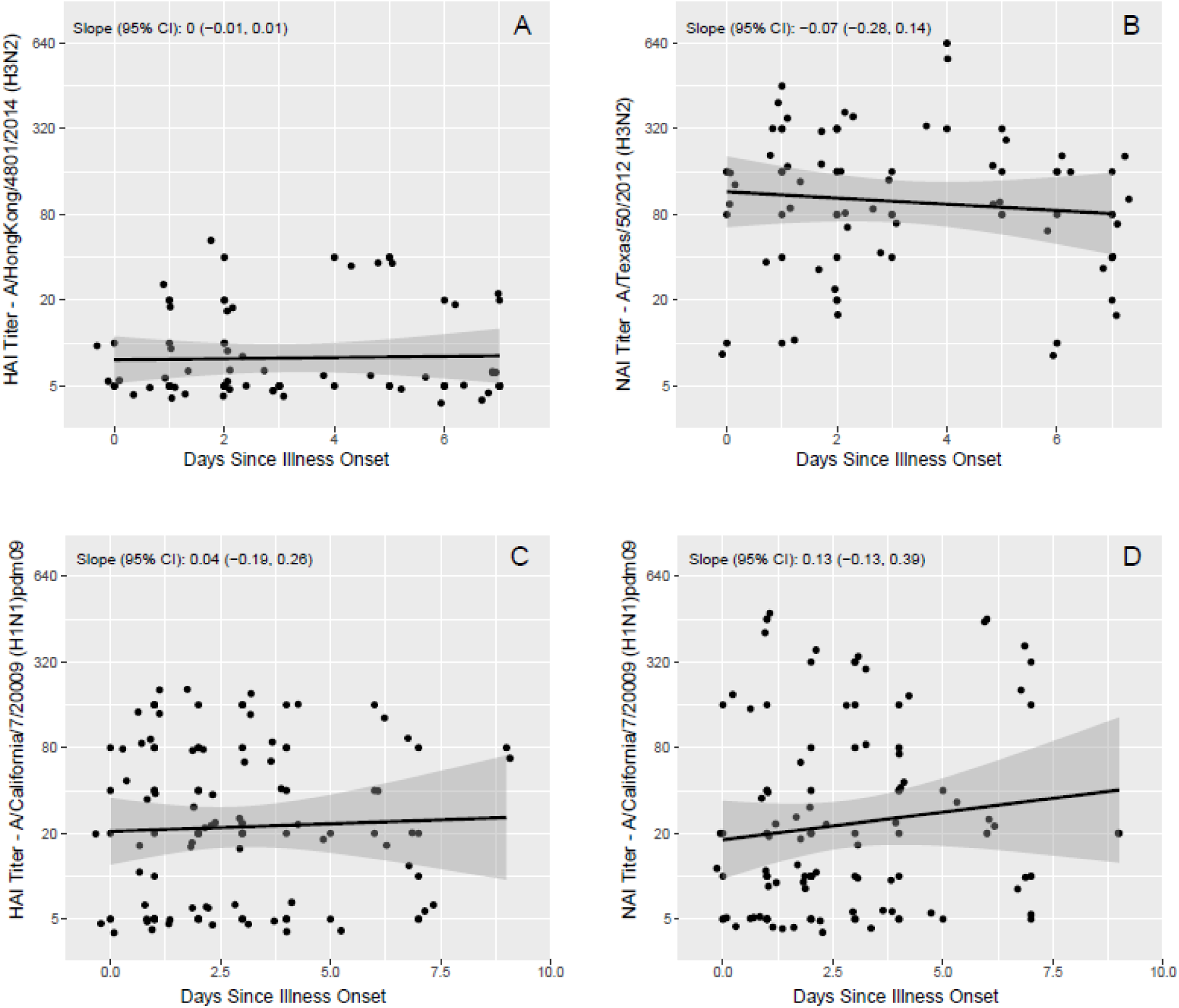
A and B: Hemagglutination-inhibition and neuraminidase-inhibition antibody titers by days from illness onset to serum specimen collection during acute illness, among influenza A(H3N2) positive cases during the 2014-2015 season. C and D: Hemagglutination-inhibition and neuraminidase-inhibition antibody titers by time from illness onset to serum specimen collection during acute illness, among influenza A(H1N1)pdm09 positive cases during the 2015-2016 season.

Among subjects enrolled in the 2014-2015 season, the overall HAI GMT was 38.8 (95% CI: 33.5, 44.8) against the A/Texas (H3N2) 3C.1 vaccine strain, and particularly low (10.4, 95% CI: 9.2, 11.8) against the A/Hong Kong (H3N2) 3C.2a that more closely resembled the circulating strain. The overall NAI GMT was 124.5 (95% CI: 109.3, 141.9) against the A/Texas (H3N2) vaccine strain (Table 1). NAI titers against A/Texas (H3N2) increased with age (*P* = 0.02), and HAI titers against A/Hong Kong (H3N2) were higher among individuals with higher frailty scores (*P*=0.04). For all viruses tested, vaccinated individuals had significantly higher A(H3N2) HAI and NAI titers than unvaccinated individuals (*P*<0.001) in these serum specimens. GMTs were higher in those not infected than those who were, but none of the differences were statistically significant (Table 1).

For the following year, the overall HAI and NAI GMTs against the A/California (H1N1)pdm09 vaccine strain were 55.9 (95% CI: 47.9, 65.3) and 50.5 (95% CI: 42.5, 59.9), respectively (Table 2). NAI GMTs against A/California (H1N1)pdm09 were higher among subjects recruited from Henry Ford Hospital (*P*=0.03), and increased with increasing age (*P*<0.001) and with increasing Charlson comorbidity index (*P*=0.01). HAI and NAI GMTs against A/California (H1N1)pdm09 were otherwise similar when compared across all other subject characteristics. For both 2015-2016 A(H1N1)pdm09 HAI and NAI, vaccinated individuals again had significantly higher GMTs than unvaccinated individuals (*P*<0.001). Influenza A(H1N1)pdm09 infected cases had significantly lower HAI and NAI GMTs against A/California (H1N1)pdm09 compared to influenza negative controls (*P*<0.001).

For the 2014-2015 season, visual examination of plots of the proportion infected by influenza A(H3N2) did not suggest a relationship between odds of infection and HAI or NAI titers against the A/Texas (H3N2) vaccine strain (Figure 2). In contrast, odds of infection did appear to decrease with increasing HAI titer against the A/Hong Kong (H3N2) strain similar to circulating 3C.2a viruses as no infections were identified among the few individuals (N=30; 10%) with titers >40. These qualitative observations were consistent with the results of logistic regression models. There was no significant association between odds of A(H3N2) infection and HAI or NAI titers against A/Texas in unadjusted, partially adjusted, or fully adjusted regression models (Table 3). For A/Hong Kong, a 2-fold increase in HAI titer was estimated to reduce odds of infection by 14% (95% CI: -19%, 48%), but this was not statistically significant.

**Figure 2.**
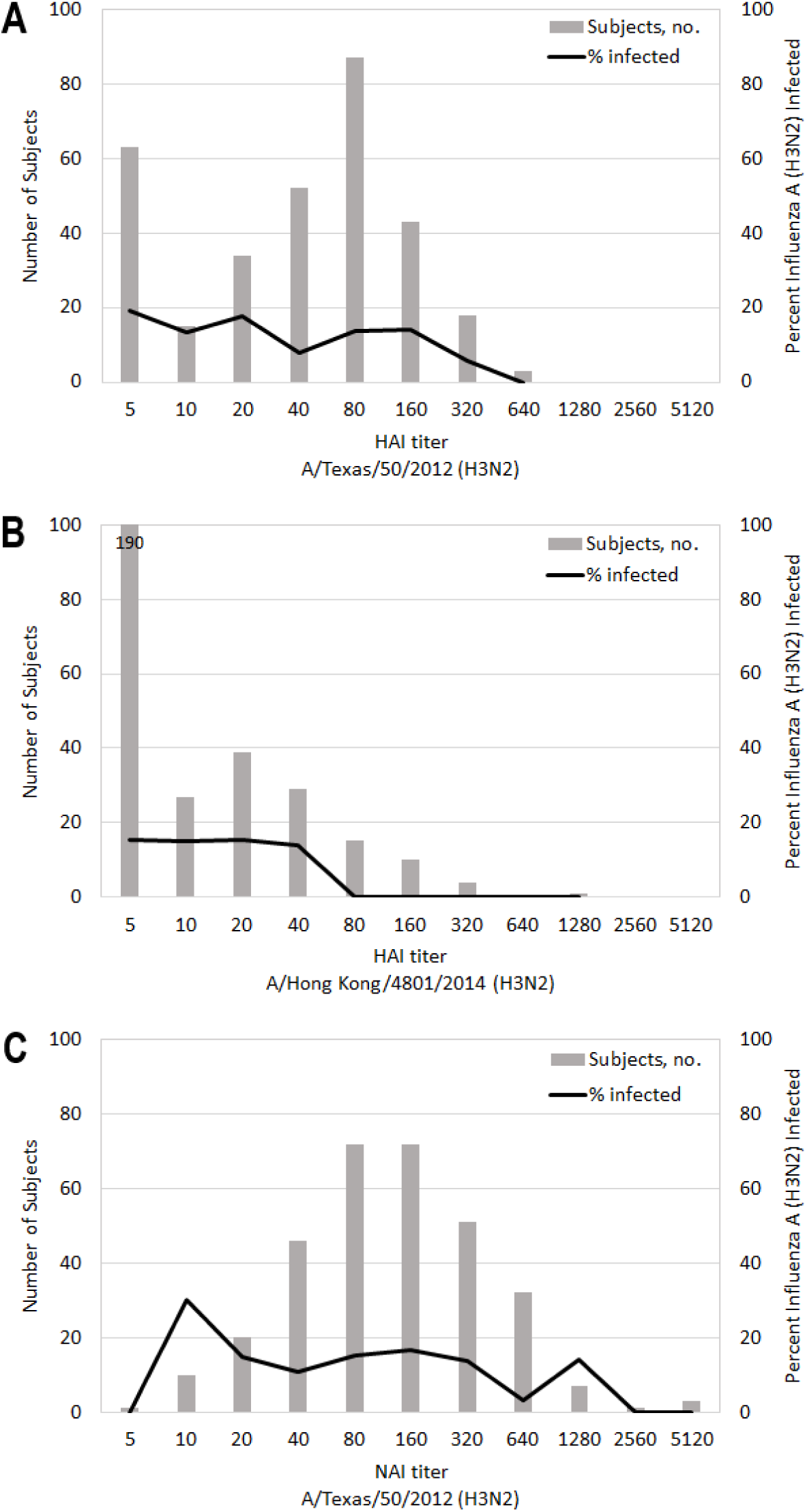
Distributions of hemagglutination-inhibition antibody titers against A) A/Texas/50/2012 (H3N2) and B) A/Hong Kong/4801/2014 (H3N2), and neuraminidase-inhibition antibody titers against C) A/Texas/50/2012 (H3N2). The solid lines in each figure represent the proportion of individuals at each titer level with RT-PCR confirmed influenza A (H3N2) infection during the 2014-2015 influenza season.

**Table 3.** Associations between hemagglutination-inhibition and neuraminidase-inhibition antibody titers and influenza infection during the 2014-2015 and 2015-2016 influenza seasons.

|  | 2014-2015 Study Year |  |  |  |  |
| --- | --- | --- | --- | --- | --- |
|  | Model 1<sup>a</sup><br>OR (95% CI) | Model 2<sup>a</sup><br>OR (95% CI) | Model 3<sup>a</sup><br>OR (95% CI) | Model 4<sup>a</sup><br>OR (95% CI) | Model 5<sup>a</sup><br>OR (95% CI) |
| HAI titer: A/Texas/50/2012 (H3N2) | 0.88 (0.75, 1.04) |  |  | 0.97 (0.78, 1.21) | 0.98 (0.78, 1.23) |
| NAI titer: A/Texas/50/2012 (H3N2) |  | 0.87 (0.72, 1.06) |  | 0.95 (0.76, 1.19) | 0.98 (0.77, 1.23) |
| HAI titer: A/Hong Kong/4801/2014 (H3N2) |  |  | 0.81 (0.61, 1.01) | 0.85 (0.61, 1.13) | 0.86 (0.62, 1.15) |
| Vaccinated for Influenza |  |  |  |  | 0.54 (0.25, 1.19) |
| Female |  |  |  |  | 1.65 (0.85, 3.26) |
| Age category (y) |  |  |  |  |  |
| 18-49 |  |  |  |  | 0.46 (0.19, 1.09) |
| 50-64 |  |  |  |  | 0.53 (0.23, 1.17) |
| ≥65 |  |  |  |  | Ref |
| Frailty score |  |  |  |  | 0.78 (0.59, 1.01) |
| CCI category |  |  |  |  |  |
| 0 |  |  |  |  | 1.25 (0.38, 3.64) |
| 1 |  |  |  |  | 1.41 (0.61, 3.15) |
| 2 |  |  |  |  | 0.78 (0.27, 1.98) |
| 3+ |  |  |  |  | Ref |
|  | 2015-2016 Study Year |  |  |  |  |
|  | Model 1<sup>a</sup><br>OR (95% CI) | Model 2<sup>a</sup><br>OR (95% CI) | Model 3<sup>a</sup><br>OR (95% CI) | Model 4<sup>a</sup><br>OR (95% CI) |  |
| HAI titer: A/California/7/2009 (H1N1)pdm09 | 0.68 (0.58, 0.78) |  | 0.73 (0.62, 0.86) | 0.78 (0.65, 0.94) |  |
| NAI titer: A/California/7/2009 (H1N1)pdm09 |  | 0.74 (0.66, 0.86) | 0.86 (0.74, 1.00) | 0.88 (0.75, 1.03) |  |
| Vaccinated for Influenza |  |  |  | 0.53 (0.27, 1.05) |  |
| Female |  |  |  | 0.91 (0.49, 1.67) |  |
| Age category (y) |  |  |  |  |  |
| 18-49 |  |  |  | 0.97 (0.42, 2.22) |  |
| 50-64 |  |  |  | 0.77 (0.35, 1.69) |  |
| ≥65 |  |  |  | Ref |  |
| Frailty score |  |  |  | 1.03 (0.82, 1.29) |  |
| CCI category |  |  |  |  |  |
| 0 |  |  |  | 3.43 (1.43, 8.26) |  |
| 1 |  |  |  | 1.30 (0.52, 3.08) |  |
| 2 |  |  |  | 2.54 (1.08, 5.81) |  |
| 3+ |  |  |  | Ref |  |

Abbreviations: 95% CI, 95% confidence interval; CCI, Charlson comorbidity index; HAI, hemagglutination-inhibition; NAI, neuraminidase-inhibition; OR, odds ratio.
<sup>a</sup> Variables with reported odds ratios and confidence intervals in a specified model column represent those included in the specified model.

In contrast, the proportion infected by A(H1N1)pdm09 in 2015-2016 appeared to decrease with increasing HAI and NAI titers against influenza A(H1N1)pdm09 (Figure 3). Consistent with this both HAI and NAI titers were significantly associated with reduced odds of infection in unadjusted and partially adjusted logistic regression models (Table 3). In fully adjusted multivariable logistic regression models, a 2-fold increase in HAI titer was associated with a 22% (95% CI: 7%, 35%) reduction in the odds of influenza A(H1N1)pdm09 infection, and a 2-fold increase in NAI titer was estimated to reduce the odds of infection by 12% (95% CI: -3%, 25%; Table 3); however, only the association between HAI titer and odds of influenza A(H1N1)pdm09 infection remained statistically significant after full covariate adjustment.

**Figure 3.**
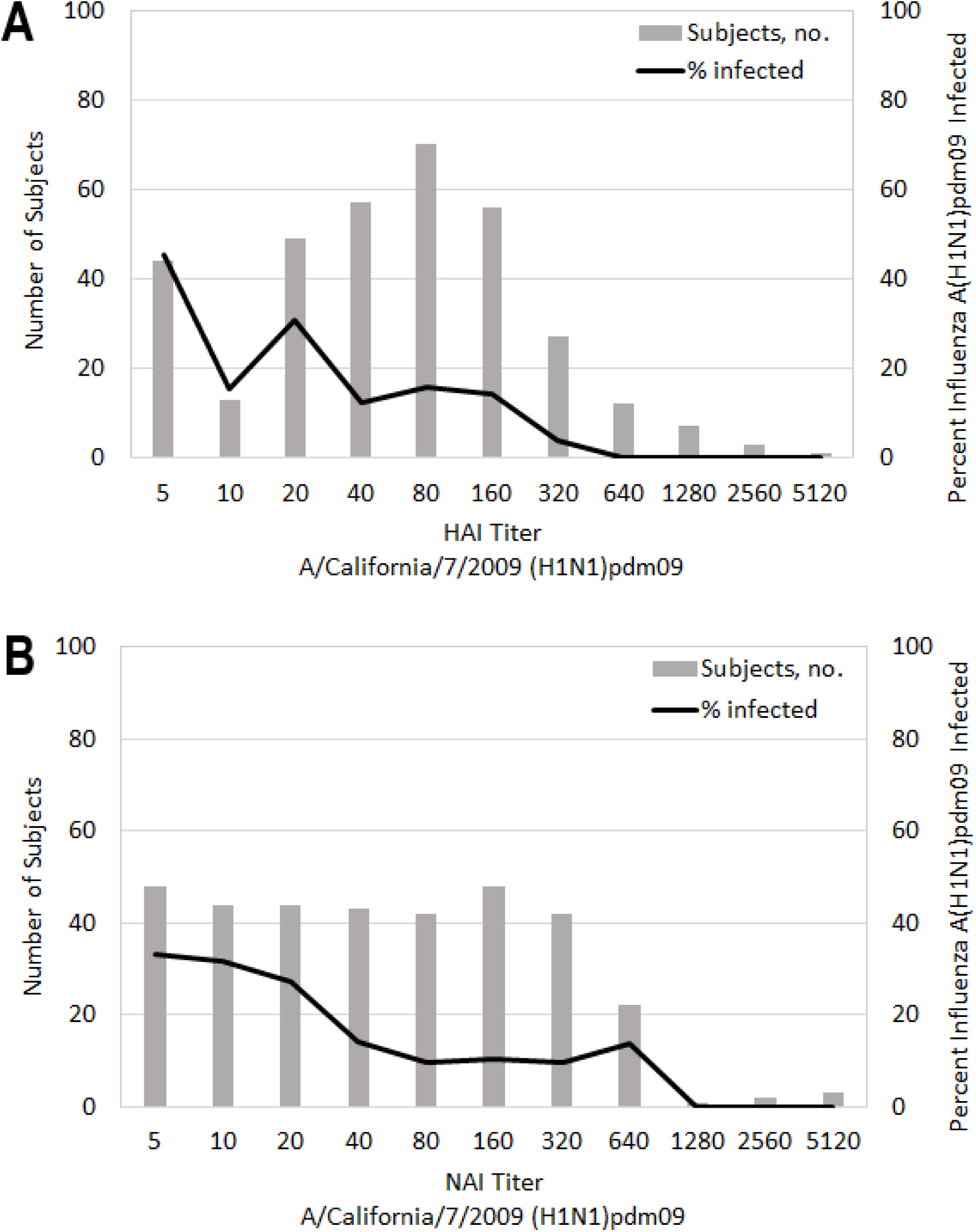
Distribution of A) hemagglutination-inhibition and B) neuraminidase-inhibition antibody titers against A/California/7/2009 (H1N1)pdm09. The solid lines in each figure represent the proportion of individuals at each titer level with RT-PCR confirmed influenza A (H1N1)pdm09 infection during the 2015-2016 influenza season.

We compared HAI and NAI antibody titers by two-year vaccination status in both study seasons (Figure 4). In each season and for each antigen, the group with the lowest titers were those unvaccinated in both seasons. In both seasons, HAI and NAI titers against all antigens did not significantly differ comparing those vaccinated only in the current season and those vaccinated in two consecutive seasons.

**Figure 4.**
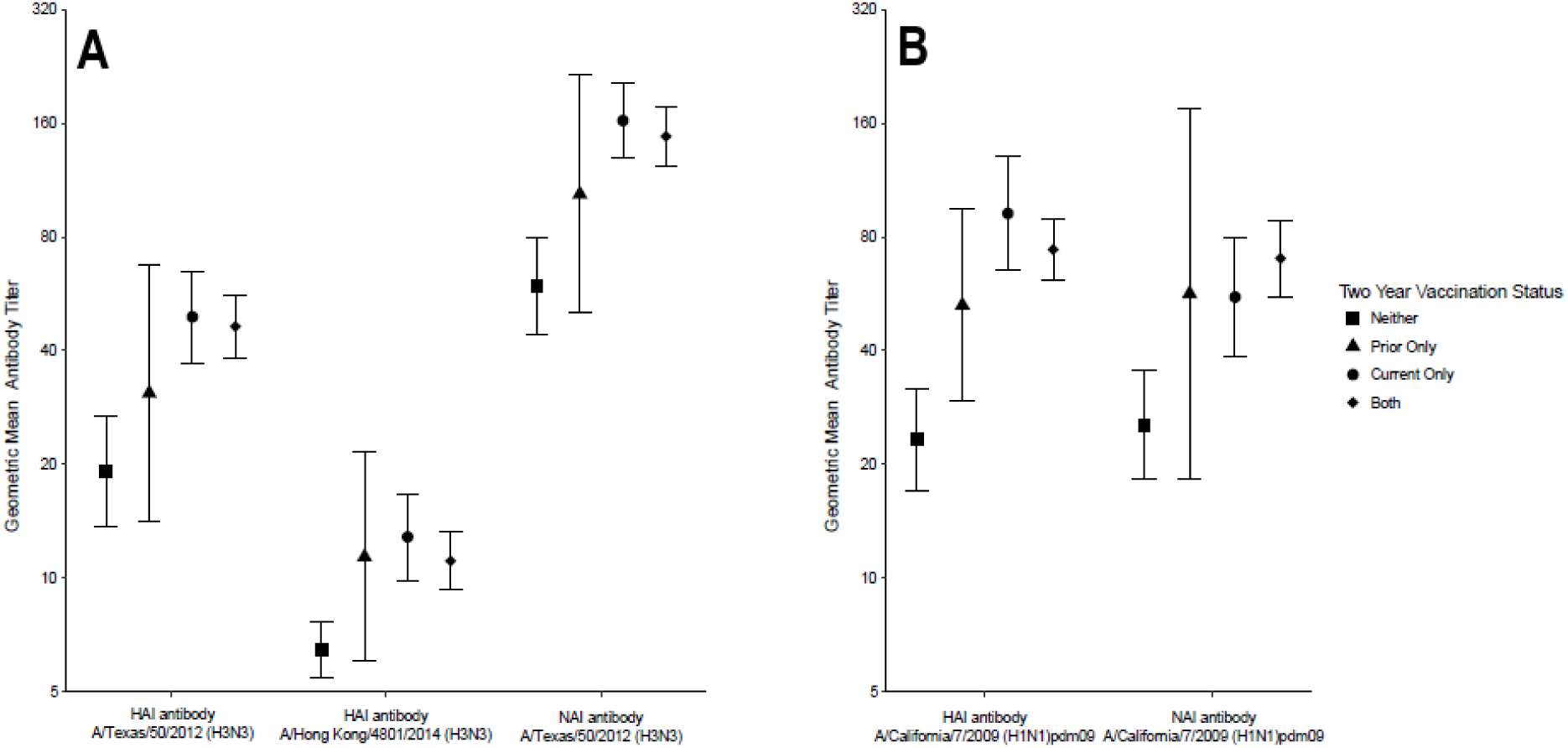
Geometric mean hemagglutination-inhibition and neuraminidase-inhibition antibody titers by two-year vaccination status in the A) 2014-2015 and B) 2015-2016 influenza seasons.

## DISCUSSION

Studies of influenza VE are now conducted annually in many parts of the word, mainly in ambulatory populations using the test negative design [4–6]. By virtue of recruiting in the ambulatory care setting it is difficult to collect adequate blood specimens from the numbers of subjects needed to evaluate serologic correlates of protection. In hospitalized individuals, with little alteration in routine practice, it is often possible to obtain residual serum specimens collected for clinical purposes. The question, in the case of study enrollment at the time of hospitalization, is whether antibodies measured in these serum specimens are indicative of pre-infection levels of susceptibility. The current results indicate that this is possible, particularly because specimens were collected shortly after acute respiratory illness onset for most patients. The fact that antibody titers were significantly higher among those who were vaccinated is also consistent with expectations for pre-infection antibody measurements.

The usefulness of these specimens is further indicated by the findings in the two influenza seasons which were divergent, but consistent with vaccine effectiveness estimates. Influenza A(H3N2) viruses which were antigenically mismatched to the A/Texas (H3N2) virus included in the vaccine circulated widely during the 2014-2015 influenza season [19,20]. Consistent with this mismatch, studies carried out in the outpatient setting observed an absence of VE in preventing infection by drifted A(H3N2) viruses [9]. In this study, HAI antibody titer against A/Texas (H3N2) did not correlate with protection against infection as expected. In contrast, there were no infections identified in the few individuals with titers ≥40 against A/Hong Kong (H3N2), which belongs to the 3C.2a genetic group that includes the majority of 2014-2015 circulating A(H3N2) viruses. Vaccinated individuals had higher titers against both viruses than unvaccinated individuals, and unlike studies carried out in ambulatory care and household settings, we found that the A/Texas (H3N2) containing vaccine was 43% effective in this hospitalized population [13].

The 2015-2016 influenza season was characterized by circulating influenza A(H1N1) viruses that were considered to be antigenically similar to the A/California (H1N1)pdm09 vaccine strain by standard HAI assays using ferret anti-sera [21]. Outpatient and inpatient based VE studies consistently estimated moderate VE in this season [17]. Consistent with this, we observed higher titers among vaccinated individuals and HAI antibody against A/California (H1N1)pdm09 was associated with protection from influenza A(H1N1)pdm09 infection in the 2015-2016 season.

Our group and others have previously demonstrated that NAI antibody is a correlate of protection against influenza infection independent of HAI antibody [22,23]. Here higher NAI titers were associated with reduced odds of infection only in the 2015-2016 season, but this effect was not statistically significant in fully adjusted models potentially reflecting low sample size. Although NA content is not standardized in influenza vaccines, it has been previously demonstrated that a substantial proportion of vaccinated individuals produce NA directed antibody in response to vaccination [22,24]. Given the potential for NA antibody to contribute to protection, more systematic monitoring of the antigenic properties of NA in circulating viruses, as is done with HA, is needed to inform appropriate targets for this relatively new assay. This will be particularly important as new approaches for vaccine production that will give broader protection are examined [12].

The primary limitation of this study is that only single serum specimens collected during acute illness were available. Although, our results support the assumption that antibody measured in these specimens are representative of levels present prior to infection, this cannot be definitively determined without paired pre-infection specimens. One recent study of antibody kinetics in patients hospitalized found that HAI titers rose rapidly after infection with >80% having HAI titers ≥40 within two weeks [25]; in our study, 89% of patients had specimens collected ≤6 days after illness onset. Similarly, vaccinated subjects had higher HAI and NAI titers against all measured influenza antigens suggesting that these antibodies were produced in response to influenza vaccination. However, because pre-vaccination serum specimens were unavailable we cannot determine how much current season vaccine contributed to the development of these antibodies.

Because enrollment in this study was cross-sectional, it is possible that individuals defined as influenza negative controls could have been infected prior to or after enrollment. This misclassification would be expected to bias estimated associations between antibody titer and odds of infection toward the null. If this bias were present, controls enrolled later in the season would be expected to have a higher probability of being infected prior to the illness for which they were enrolled, and therefore, higher antibody titers than those enrolled earlier in the season. We did not observe such a relationship, and therefore, would expect any misclassification of infection status to be minimal. Because this study was carried out in only two hospitals in Southeast Michigan, the generalizability of the results to other populations is unclear; however, the results are generally consistent with larger, national studies of influenza vaccine effectiveness. The results of this study should also be interpreted in the context of limited sample size.

The case-test negative design has allowed for efficient annual estimation of the effectiveness of influenza vaccines. These studies have been critical in informing public health policy by identifying issues with current influenza vaccines such as the poor performance of live-attenuated influenza vaccines in recent years [17,26]. More generally, these studies have consistently reported low VE against influenza A(H3N2) viruses even in seasons when the hemagglutinins of vaccine strain viruses are considered well matched to those that circulate [7]. Pairing these studies with immunologic evaluations, though difficult, is currently a major unmet need for interpreting annual VE estimates. The results of this study support the utility of using serum specimens collected during acute illness to assess correlates of protection against influenza infection to support hospital-based studies of influenza VE. Implementation of similar studies more broadly would provide an opportunity to understand the successes and shortcomings of current influenza vaccines.

## Supplementary Materials

### Frailty Score

During the enrollment interview, each participant was asked five questions comprising a frailty short interview based on the components of the frailty phenotype defined by Fried et al. [27]. Participants were asked the following questions:

“In the last year, before this current illness, have you lost more than 10 pounds unintentionally (i.e., not due to dieting or exercise)?” [YES;NO]; adapted from [27].
“In the last month, before this current illness, have you had too little energy to do the things you wanted to do?” [YES;NO]; adapted from [28].
“Before this current illness, how difficult was it for you to lift or carry something as heavy as 10 pounds, such as a full bag of groceries, by yourself, and without using any special equipment?” [NO DIFFICULTY; A LITTLE…; SOME…; A LOT…; UNABLE TO DO]; adapted from [29].
“Before this current illness, did you have difficulty walking 100 yards (around the size of a football field) because of a health problem?” [NO DIFFICULTY; A LITTLE…; SOME…; A LOT…; UNABLE TO DO]; adapted from [28].
“Before this current illness, how often did you engage in activities that require a low or moderate level of energy such as gardening, cleaning the car, or going for a walk?” [>ONCE PER WEEK; ONCE PER WEEK; 1-3 TIMES PER MONTH; HARDLY EVER/NEVER]; adapted from [28].

Responses to the last three questions were recoded as dichotomous variables with at least some difficulty or engaging in physical less than once per week indicating positive responses indicative of frailty. The responses to each element of the five elements were summed to create a frailty score (values 0 to 5) which was included as a continuous covariate in multivariable regression models.

### Hemagglutination-inhibition assay

The hemagglutination-inhibition assay (HAI) assays were performed to measure antibodies that block hemagglutinin (HA) receptor binding. Prior to HAI testing, all sera were treated overnight with receptor destroying enzyme and heat inactivated to prevent nonspecific inhibition; sera were also adsorbed with red blood cells to remove nonspecific agglutinins. Serial 2-fold dilutions (with an initial dilution of 1:10) were prepared for each serum specimen in 96-well microtiter plates. The serum dilutions were then incubated with standardized concentrations (4 HA units per 25 μL) of monovalent IIV subunit material (provided by Sanofi-Pasteur) representing the A/Texas/50/2012 (H3N2) virus present in the 2014-2015 Northern Hemisphere influenza vaccine and the A/Hong Kong /4801/2014 (H3N2) virus present in the 2016-2017 Northern Hemisphere influenza vaccine and belonging to the 3C.2a genetic group. Turkey red blood cells were added to wells and allowed to settle. The strain-specific HAI antibody titers at each time point for each individual were calculated as the reciprocal (eg, 40) of the highest dilution of sera (eg, 1:40) that inhibited hemagglutination. HAI titers below the limits of detection (ie, <10) were denoted as half of the threshold detection value (ie, 5); titers greater than the upper test value (ie, 5120) were denoted as having twice that value (ie, 10240).

### Enzyme-linked lectin assay

Neuraminidase-inhibition (NAI) antibody titers were measured by enzyme-linked lectin assay (ELLA) [30]. This assay used a reassortant influenza viruses with a mismatched HA (H6 subtype), to avoid interference by HA-specific antibodies, and with the neuraminidase (NA) antigen representing either the A/Texas/50/2012 (H3N2) virus present in the 2014-2015 Northern Hemisphere influenza vaccine or the A/California/7/2009 (H1N1)pdm09 virus present in the 2015-2016 Northern Hemisphere influenza vaccine. Serum specimens were heat inactivated, and serial 2-fold dilutions (with an initial dilution of 1:10) were incubated with virus and then added to 96-well microtiter plates coated with fetuin in duplicate. A 25mM MES buffer is used throughout the assay as diluent and for plate washing. A final solution of 20mM CaCl2 and 1% BSA are added to the diluents with a pH of 6.5. Following incubation, peroxidase-labeled peanut agglutinin (the lectin) and, later, peroxidase substrate were added to detect enzymatic cleavage of fetuin by viral NA, and the reaction optical density was measured with a microplate reader. The percentage inhibition of NA enzymatic activity at each serum dilution was calculated by comparison with values from virus control wells (virus but no serum); endpoint NAI titers were calculated as the reciprocal of the highest dilution with at least 50% inhibition.

---

**Supplementary Table 1.** Comparison of subject characteristics by serum specimen availability among adults hospitalized for acute respiratory illness during the 2014-2015 influenza season.

| Characteristic | University of Michigan Hospital |  |  |
| --- | --- | --- | --- |
|  | Total Participants Enrolled No. | Specimens Retrieved No. (%) | P-value<sup>a</sup> |
| Overall | 341 | 315 (92) | - |
| Sex |  |  | 0.65 |
| Female | 169 | 155 (92) |  |
| Male | 172 | 160 (93) |  |
| Age category (y) |  |  | 0.74 |
| 18-49 | 90 | 84 (93) |  |
| 50-64 | 108 | 98 (91) |  |
| ≥65 | 143 | 133 (93) |  |
| Race/ethnicity<sup>b</sup> |  |  | 0.96 |
| White (not Hispanic) | 262 | 242 (92) |  |
| Black | 47 | 43 (91) |  |
| Other | 30 | 28 (93) |  |
| Frailty score (range 0-5) |  |  | 0.49 |
| 0-2 | 243 | 226 (93) |  |
| 3-5 | 98 | 89 (91) |  |
| Charlson comorbidity index category |  |  | 0.25 |
| 0 | 34 | 33 (97) |  |
| 1 | 81 | 71 (88) |  |
| 2 | 56 | 53 (95) |  |
| 3+ | 170 | 158 (93) |  |
| Influenza vaccination status |  |  | 0.12 |
| Vaccinated | 253 | 237 (94) |  |
| Unvaccinated | 88 | 78 (89) |  |

<sup>a</sup> P-values are calculated from chi-square tests.
<sup>b</sup> 2 participants with serum specimens had missing Race/ethnicity data

**Supplementary Table 2.** Comparison of subject characteristics by serum specimen availability among adults hospitalized for acute respiratory illness during the 2015-2016 influenza season.

| Characteristic | University of Michigan Hospital |  |  | Henry Ford Hospital |  |  |
| --- | --- | --- | --- | --- | --- | --- |
|  | Total Participants Enrolled No. | Specimens Retrieved No. (%) | P-value<sup>a</sup> | Total Participants Enrolled No. | Specimens Retrieved No. (%) | P-value<sup>a</sup> |
| Overall | 257 | 243 (95) | - | 184 | 96 (52) | - |
| Sex |  |  | 0.47 |  |  | 0.53 |
| Female | 134 | 128 (96) |  | 117 | 59 (50) |  |
| Male | 123 | 115 (94) |  | 67 | 37 (55) |  |
| Age category (y) |  |  | 0.27 |  |  | 0.40 |
| 18-49 | 86 | 79 (92) |  | 60 | 27 (45) |  |
| 50-64 | 81 | 79 (98) |  | 77 | 43 (56) |  |
| ≥65 | 90 | 85 (94) |  | 47 | 26 (55) |  |
| Race/ethnicity<sup>b</sup> |  |  | 0.66 |  |  | 0.03 |
| White (not Hispanic) | 182 | 172 (95) |  | 37 | 26 (70) |  |
| Black | 35 | 34 (97) |  | 128 | 61 (48) |  |
| Other | 39 | 36 (92) |  | 14 | 5 (36) |  |
| Frailty score (range 0-5) |  |  | 0.83 |  |  | 0.28 |
| 0-2 | 177 | 167 (94) |  | 131 | 65 (50) |  |
| 3-5 | 80 | 76 (95) |  | 53 | 31 (58) |  |
| Charlson comorbidity index category |  |  | 0.30 |  |  | 0.02 |
| 0 | 29 | 27 (93) |  | 36 | 14 (39) |  |
| 1 | 40 | 36 (90) |  | 47 | 19 (40) |  |
| 2 | 36 | 33 (92) |  | 17 | 12 (71) |  |
| 3+ | 152 | 147 (97) |  | 84 | 51 (61) |  |
| Influenza vaccination status |  |  | 0.16 |  |  | 0.46 |
| Vaccinated | 199 | 186 (93) |  | 93 | 51 (55) |  |
| Unvaccinated | 58 | 57 (98) |  | 91 | 45 (49) |  |

<sup>a</sup> P-values are calculated from chi-square tests.
<sup>b</sup> 5 participants with serum specimens had missing Race/ethnicity data

**Supplementary Figure 1.**
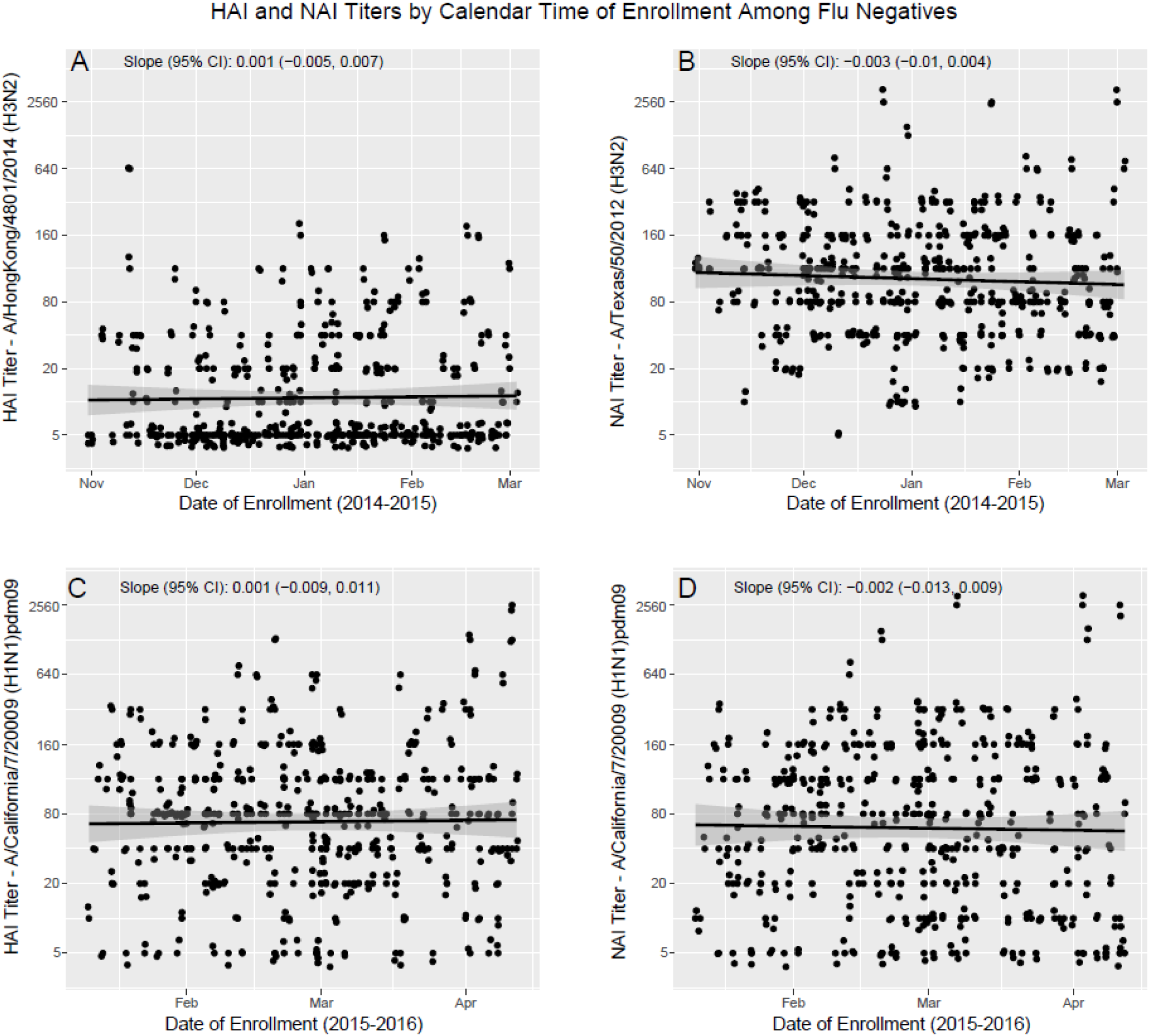

## Acknowledgements

We thank Jin Gao and Laura Couzens for technical support and are indebted to St Jude Children’s Research Hospital for plasmids that were used to generate reassortant influenza viruses.

## Notes

**Financial Support:** This work was supported by the Centers for Disease Control and Prevention (U01 IP000974, U01 IP000474).

**Potential conflicts of interest:** E.T.M has received grant support from Merck and Pfizer for work unrelated to this report. L.E.L. has received grant support from AstraZeneca, Merck, Pfizer, eMaxHealth Inc., Policy Analysis Inc., Analytica Inc. and Xcenda Inc. for work unrelated to this report. A.S.M. has received grant support from Sanofi Pasteur and consultancy fees from Sanofi, GSK and Novavax for work unrelated to this report. All authors reported no potential conflicts.

